# PHF2 regulates homology-directed DNA repair by controlling the resection of DNA double strand breaks

**DOI:** 10.1101/782490

**Authors:** Ignacio Alonso-de Vega, M. Cristina Paz-Cabrera, Wouter W. Wiegant, Cintia Checa-Rodríguez, Pablo Huertas, Raimundo Freire, Haico van Attikum, Veronique A.J. Smits

## Abstract

Post-translational histone modifications and chromatin remodelling play a critical role in the mechanisms controlling the integrity of the genome. Here we identify histone lysine demethylase PHF2 as a novel regulator of the DNA damage response by regulating the balance between DNA damage-induced focus formation by 53BP1 and BRCA1, critical factors in the pathway choice for DNA double strand break repair. PHF2 knock down leads to impaired BRCA1 focus formation and delays the resolution of 53BP1 foci. Moreover, irradiation-induced RPA phosphorylation and focus formation, as well as localization of CtIP, required for DNA end resection, to sites of DNA lesions are affected by depletion of PHF2. These results are indicative of a defective resection of double strand breaks and thereby an impaired homologous recombination upon PHF2 depletion. In accordance with these data, Rad51 focus formation and homology-directed double strand break repair is inhibited in cells depleted for PHF2. Importantly, we demonstrate that PHF2 knock down decreases CtIP and BRCA1 protein and mRNA levels and cells depleted of PHF2 display genome instability and are sensitive to the inhibition of PARP. Together these results demonstrate that PHF2 promotes DNA repair by homologous recombination by controlling CtIP-dependent resection of double strand breaks.

## INTRODUCTION

The DNA damage response (DDR), which detects, signals and repairs DNA lesions, is essential in the maintenance of genome integrity and functions as a first defence in the early stages of cancer development (1). Among the different types of DNA lesions, DNA double strand breaks (DSBs) are particularly hazardous to the cell as, if not repaired adequately, they can lead to chromosomal rearrangements (2).

Mammalian cells have developed different pathways to repair DSBs. Non-homologous end joining (NHEJ) is a fast, and efficient, but also error-prone pathway in which the broken DNA-ends are directly ligated. On the other hand, during homologous recombination (HR), the sister chromatid is used as a repair template and thereby results in more accurate repair. Finally, the less efficient alternative non-homologous end joining (Alt-NHEJ), also called microhomology-mediated end joining (MMEJ), uses short microhomologous sequences during the alignment of broken ends (3). Several factors are important in DSB repair pathway choice, in which the availability of homologous sequence, and therefore the cell cycle stage, plays a critical role. Two DDR proteins that have an important influence on this decision are 53BP1 and BRCA1, that together control DNA end resection, the degradation of the DNA end in the 5’ to 3’ direction, resulting in singe-stranded (ss) DNA that is critical for DSB repair by HR (4). Whereas 53BP1, together with its partner RIF1 and the Shieldin complex (5, 6), blocks DNA end-resection and thus stimulates repair through NHEJ, BRCA1 promotes DNA end-resection and the removal of 53BP1 from sites of DNA damage, thereby switching repair from NHEJ to HR (7, 8). DNA end resection is initiated by endonuclease CtIP, in cooperation with the Mre11/Rad50/Nbs1 (MRN) complex, and thereafter extended by the EXO1 and DNA2 nucleases (9). The resulting ssDNA is protected by immediate coating with the RPA1-3 complex, which is replaced by the Rad51 protein that then mediates strand invasion (10).

Efficient DNA repair requires the correct and timely coordination of a multitude of signalling events, in which posttranslational modifications play a critical role. As DNA lesions occur and are repaired in the context of chromatin, it is not surprising that chromatin modifications impact on this process (11). One of the earliest modifications upon the induction of a DSB is the phosphorylation of histone H2AX at serine 139 (named γH2AX) by the central DDR kinase ATM, on either side of the lesion (12). H2AX phosphorylation triggers the initiation of protein ubiquitination by the RNF8 E3 ubiquitin ligase, which is recruited to DSBs via binding to MDC1, a direct reader of γH2AX (13). Ubiquitination of histones and other proteins by RNF8 and RNF168, another E3 ligase that is subsequently bound, serves to recruit additional proteins, among which are 53BP1 and BRCA1 (14, 15). Also, the modification of histone and non-histone proteins by methylation and acetylation are involved in the regulation of DNA repair. For example, the recruitment of 53BP1 depends on the methylation of H4K20, the RNF8-dependent degradation of competing H4K20me readers, deacetylation of H4K16 and RNF168-mediated H2AK15 ubiquitination (16). In addition to promoting the direct recruitment of repair proteins, chromatin modifications can physically facilitate the accessibility of regulatory proteins to the lesion. An example is lysine acetylation, which opens up the chromatin (17). Finally, histone methylation regulates gene expression, by recruiting proteins involved or by inhibiting the binding of transcription factors to DNA. Lysine methylation of histone H3 and H4 is associated to both transcriptional activation and repression, depending on the methylation site (18).

Interestingly, defects in the regulation of chromatin modifying enzymes were described to be linked to genome instability and tumor development (19). As additional chromatin modifications and regulators are likely yet to be discovered to be involved in the DDR and in DNA repair in particular, we set out to identify novel regulators of chromatin modifications that regulate the balance between NHEJ and HR and identified lysine-specific demethylase PHF2 as a regulator of homology-directed DNA repair by controlling CtIP and BRCA1.

## MATERIALS AND METHODS

### Cell culture and treatments

293T, U2OS and HeLa cells were grown in DMEM supplemented with 10% FBS and penicillin-streptomycin. U2OS DR-GFP and U2OS SA-GFP reporter stable cell lines were maintained in medium supplemented with 1 µg/ml puromycin. Rap80 knock out U2OS cells were kindly provided by Dr. Daniel Durocher (Lunenfeld-Tanenbaum Research Institute, Mount Sinai Hospital, Toronto, Canada) and have been validated previously (20).

Cells were treated with camptothecin (CPT, 2 μM), hydroxyurea (HU, 10 mM), ultra violet light (UV, 40 J/m2), etoposide (ETP, 20 μM) or ionizing radiation by a CellRad (Faxitron) and harvested one hour later, unless stated otherwise.

### siRNA oligos, plasmids and transfection

siRNA oligonucleotides (Sigma) were transfected into cells using Lipofectamine RNAiMax (Invitrogen). Sequences of oligonucleotides were as follows:

Luc CGUACGCGGAAUACUUCGA

Non-Target UGGUUUACAUGUCGACUAA

PHF2#1 GCUGGAAAUUCGAGAGCAA

PHF2#2 GCUAGAGAAGUCGCCUCUA

PHF2#3 CCACUUUAAGGACAGCCUU

CtIP GCUAAAACAGGAACGAAUC

BRCA1 AAGGAACCUGUCUCCACAAAG

Plasmid DNA was transfected into cells using Polyethylenimine (Sigma Aldrich) or Lipofectamine 2000 (Invitrogen).

p3FLAG murine PHF2 was kindly provided by Dr. Xiaobing Shi (Department of Epigenetics and Molecular Carcinogenesis, The University of Texas M. D. Anderson Cancer Center, Houston, USA). Three silent mutations (in capital, gctggaGatCAgGgagcaa) were introduced in the Flag-PHF2 plasmid using the QuikChange Site-Directed Mutagenesis Kit (Agilent Technologies) to make it resistant to siRNA oligonucleotide PHF2#1 (Flag-PHF2*). GFP-CtIP and mCherry-NBS1 have been previously described (21, 22).

### Antibodies and western blot

Antibodies obtained from commercial sources were as following: β-actin and Histone H3 from Genscript, Ku86 (C-20) and p53 (DO-1) from Santa Cruz Biotechnology, 53BP1 (Ab172580) and NBS1 (Ab175800) from Abcam, pSer139-H2AX (clone: JWB301), BRCA1 (clone MS110) from Merck-Millipore, pSer345-CHK1 and PHF2 from Cell Signalling, PHF2 and pSer4/8-RPA2 from Bethyl, RPA2 from Novus Biologicals, Rad51 by Invitrogen and CtIP from Active Motif.

The antibodies against CtIP and Mre11 were generated by injecting rabbits with a His-tagged antigen (amino acids 150-500 and 182-480, respectively) that was obtained by expression in bacteria and purified with a Ni-NTA resin (Qiagen) following manufacturers recommendations.

Cell were lysed in Laemmli sample buffer. Lysates containing equal amounts of protein, measured by the BCA method (Thermo Scientific), were subjected to SDS-PAGE. The chemiluminescent images were obtained using the ImageQuant LAS 4000 mini (GE Healthcare).

### Comet assay

Single Cell Gel Electrophoresis (SCGE) was carried out using the CometAssay® ES II kit (Trevigen) according to the manufacturer’s instructions. Images were taken using a Zeiss Cell Observer fluorescent microscope and the tail moment of at least 50 cells per experiment was analysed with the TriTek CometScore software.

### RT-PCR

RNA was isolated using the RiboZol Extraction Reagent (VWR) according to the manufacturer’s instructions. cDNA synthesis and PCR amplification were carried out in the same tube using the qScript One-Step SYBR Green qRT-PCR Kit (Quantabio) and a LightCycler480 II (Roche). All reactions were performed in triplicate. Transcript levels were normalized in parallel with test genes GAPDH. The primers used were the following:

BRCA1-F: ACCTTGGAACTGTGAGAACTCT

BRCA1-R: TCTTGATCTCCCACACTGCAATA

CtIP-F: CAGGAACGAATCTTAGATGCACA

CtIP-R: GCCTGCTCTTAACCGATCTTCT

GAPDH-F: GGAGCGAGATCCCTCCAAAAT

GAPDH-R: GGCTGTTGTCATACTTCTCATGG

Mre11-F: ATGCAGTCAGAGGAAATGATACG

Mre11-R: CAGGCCGATCACCCATACAAT

PHF2-F: TTCTCTGACACCCGAATGTCC

PHF2-R: CCTTCACGCAGATTAGGCAGT

Rad51-F: CGAGCGTTCAACACAGACCA

Rad51-R: GTGGCACTGTCTACAATAAGCA

RPA2-F: GCACCTTCTCAAGCCGAAAAG

RPA2-R: CCCCACAATAGTGACCTGTGAAA

### DR-GFP and SA-GFP assays

U2OS cells bearing a single copy integration of the reporters DR-GFP (23) or SA-GFP (24) were used to analyse the different recombination pathways. DR-GFP contains two differentially mutated GFP cassettes, one of which contains the I-SceI restriction site. The SA-GFP reporter contains two separated fragments of the GFP gene, one of which containing an I-SceI restriction site. After DSB induction by I-SceI expression, a functional GFP cassette is generated by homologous recombination or SSA-mediated repair, respectively. In both cases, the resulting GFP-positive cells are analysed by flow cytometry.

Cells were plated in 6-well plates in duplicate and transfected with the indicated siRNA oligonucleotide. The next day, cells were infected with lentiviral particles containing I-SceI–BFP expression construct at MOI 10 using 8 μg/ml polybrene. 48 h later, cells were collected by trypsinization and fixed with 4% paraformaldehyde for 20 min. Samples were analysed with a BD FACSAria with the BD FACSDiva Software v5.0.3.

The number of green cells from at least 10,000 events positives for blue fluorescence (infected with the I-SceI–BFP construct) was scored. The average of both duplicates was calculated for each sample and normalized to siRNA control. At least three independent experiments were carried out for each condition and the average and standard deviation of the three experiments represented.

### Chromatin fractionation

Biochemical fractionation of cells was performed as previously described (25, 26). Soluble cytoplasmic and soluble nuclear fractions were pooled to one soluble fraction.

### Immunofluorescence

For immunostaining, cells were fixed in 4% paraformaldehyde for 15 min and then permeabilized with PBS+0.2% Triton X-100 for 10 min at RT. Samples were blocked in PBS+0.5% FCS, immunostained with antibodies as indicated and mounted with DAPI. For RPA2, BRCA1 and Rad51 focus formation, cells were pre-extracted (20mM Hepes pH 8, 20mM NaCl, 5mM MgCl_2_, 0.5% NP40) for 5 min at 4°C before fixation.

Images of cells were taken using a Zeiss Cell Observer fluorescent microscope equipped with a 63x NA 1.3 water immersion objective and ZEN imaging software. The number of foci was evaluated using the ImageJ software. In all instances, more than 100 cells were analysed for each point and error bars on graphs represent the standard deviation of three independent experiments. Cells with more than 10 foci were scored as positive.

### Laser micro-irradiation

Multiphoton laser micro-irradiation was essentially performed as described previously (27). Cells, grown on coverslips, were placed in a Chamlide CMB magnetic chamber and the medium was replaced by CO_2_-independent Leibovitz’s L15 medium supplemented with 10% FCS and penicillin-streptomycin. Laser micro-irradiation was carried out on a Leica SP5 confocal microscope equipped with an environmental chamber set to 37°C. DSB-containing tracks (1.5 μm width) were generated with a Mira mode-locked titanium-sapphire (Ti:Sapphire) laser (*λ* = 800 nm, pulse length = 200 fs, repetition rate = 76 MHz, output power = 80 mW) using a UV-transmitting 63× 1.4 NA oil immersion objective (HCX PL APO; Leica). Confocal images were recorded before and after laser irradiation at 5 or 10 s time intervals over a period of 5–10 min.

To examine accumulation of Flag-PHF2, a different field was irradiated every minute for 20 minutes, after which the cells were fixed for immunofluorescence.

### FokI assays

U2OS 2-6-3 cells expressing inducible FokI-mCherry-LacR (28) were treated with 300 nM 4-OHT and 1 M Shield-I for 5 h for inducing stabilization and nuclear localization of the expressed product. Subsequently, cells were fixed with PFA and immunostained with the indicated antibodies as described above.

### Flow Cytometry

For cell cycle analysis, cells were fixed in 70% ethanol at 4°C o/n. After fixation, cells were washed with PBS, and the DNA was stained with propidium iodide (PI). Cells were analysed using a Macsquant Analyzer with Macsquantify software (Miltenyi).

### Clonogenic survival

To determine cellular sensitivity to DNA damaging agents, HeLa cells were transfected with the corresponding siRNA oligonucleotides. 48h later, 500 cells were seeded in 6 well dishes and treated with the indicated concentrations of Olaparib (Cayman Chemical). Following 7-10 days in culture, cells were fixed, stained and colonies were counted. Triplicate cultures were scored for each treatment. Shown is the relative survival as compared to the undamaged control and the error bars present the standard error of three independent experiments.

### Statistical analysis and reproducibility

A two-tailed Student’s t-test was used to determine whether the difference between the means of two sets of values was significant. *P<0.1 **P<0.01, ***P<0.001, ****P<0.0001. P<0.05 was considered statistically significant. NS = not significant. Unless stated otherwise, representative experiments are shown out of at least two independent ones and depicted is the mean ± standard deviation. Additional details are listed in the individual figure legends.

## RESULTS

The demethylase PHF2 controls the DNA damage response

To identify novel factors that regulate the balance between NHEJ and HR, we analysed focus formation of 53BP1 and BRCA1, that together control the choice between these two DSB repair pathways, in response to ionizing radiation (IR) in cells depleted for individual enzymes involved in post-translation modifications of histones by siRNA. Modulation of the expression level of PHF2/KDM7C/JHDM1E, a lysine-specific histone demethylase hereafter called PHF2, changed the dynamics of 53BP1 focus formation in response to IR. Whereas irradiating U2OS cells triggered efficient 53BP1 focus formation, the number of cells with 53BP1 foci and the number of 53BP1 foci per cell stayed high at later time points (4-7h) in cells depleted for PHF2 by siRNA whereas at these time points 53BP1 foci decreased again in control transfected populations (Fig. 1A and S1A). The effect of PHF2 depletion on 53BP1 focus resolution was the same as that of downregulation of CtIP, which served as a control for negatively regulating HR by preventing the initiation of DNA resection (Fig. 1A). In contrast, overexpression of Flag-PHF2 prevented IR-induced focus formation of 53BP1 (Fig. S1B). To examine the effect of modulating PHF2 levels on HR, we monitored the recruitment of BRCA1 in Rap80 knock out U2OS cells, in which Rap80-mediated recruitment of BRCA1, subsequent BRCA1-A complex formation and suppression of HR are prevented (27, 29). Whereas treating these cells with IR led to efficient focus formation of BRCA1 that increased in time, depletion of PHF2 by siRNA impairs IR-induced BRCA1 foci (Fig. 1B). Given the opposite effect of PHF2 depletion on 53BP1 versus BRCA1 IR-induced focus formation, these data suggest that PHF2 regulates the DSB response in a mechanism that is beneficial for HR and therefore detrimental for NHEJ.

**Figure 1.**
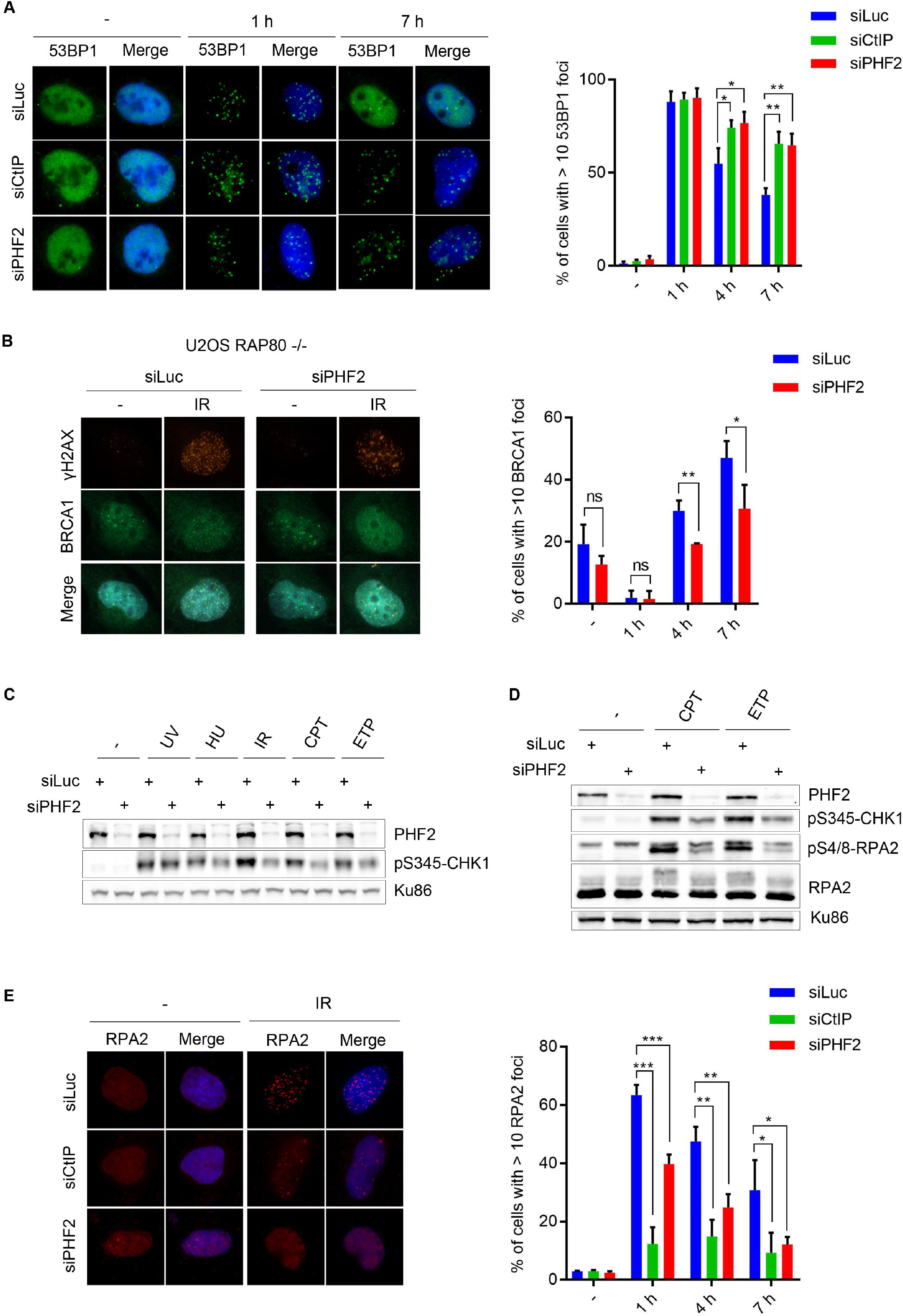
PHF2 knock down alters 53BP1 and BRCA1 focus dynamics in response to IR. (**A**) U2OS cells were depleted for Luciferase (Luc), CtIP or PHF2 by siRNA. After 48 h, cells were treated with IR (10 Gy) and fixed after 1, 4 or 7 hours. 53BP1 focus formation was analysed by immunofluorescence. Left panel: representative images. Right panel: quantification of three independent experiments with each at least 100 cells. (**B**) U2OS Rap80 knock out cells were depleted for Luc or PHF2. Cells were treated with IR (10 Gy), fixed at the indicated time points and γH2AX (positive control for DNA damage induction) and BRCA1 focus formation was analysed as in (A). (**C**) U2OS cells depleted for Luc or PHF2 were subjected to different DNA damaging agents and lysed after 1 h. Extracts were analysed by western blot using the indicated antibodies. (**D**) U2OS cells were transfected as in (C), treated with CPT or ETP and analysed by western blot with the indicated antibodies. (**E**) U2OS cells were depleted for Luc, CtIP or PHF2 by siRNA. Cells were treated with IR (3 Gy), fixed at the indicated time points and RPA2 focus formation was analysed as in (A).

### Depletion of PHF2 affects DSB resection and DNA repair by homologous recombination

We continued to study what process in the DNA damage response is affected by PHF2. To this end, we first examined DNA damage-induced checkpoint activation by exposing U2OS cells to different types of DNA damage and monitoring the phosphorylation of CHK1, a critical effector kinase of this response (30). Although DNA damaging agents triggered efficient CHK1 phosphorylation in control depleted cells, PHF2 depletion reduced the DNA damage-induced phosphorylation of CHK1, especially after IR, camptothecin and etoposide (Fig. 1C). These treatments result in a more efficient DSB formation compared to UV and HU, which only resulted in a moderate effect, substantiating a possible role for PHF2 in the DSB response. In addition, these treatments need DNA end resection to generate ssDNA, subsequently covered by the RPA1-3 complex, that triggers the activation of ATR and phosphorylation of its substrates such as CHK1 (31, 32). We therefore examined if PHF2 depletion affected DNA end resection, by studying RPA2 phosphorylation using western blot analysis (33). Downregulation of PHF2 led to lower levels of RPA2 phosphorylation on Ser4/8 and reduced levels of total RPA2 phosphorylation, as demonstrated by a lower mobility shift using an antibody against total RPA2, in response to camptothecin and etoposide, as compared to control transfected cells (Fig. 1D). To corroborate these findings, we also measured IR-induced focus formation of RPA2 by immunofluorescence. As previously published, depletion of CtIP completely inhibited focus formation of RPA2 (Fig. 1E). Notably, although less pronounced when compared to that after CtIP knock down, the knock down of PHF2 also significantly reduced IR-induced focus formation of RPA (Fig. 1E).

DSB resection is initiated by CtIP, together with the MRN complex. Given the effect of PHF2 on DNA end resection, we wondered if PHF2 functions at the level of CtIP. We therefore monitored the recruitment of GFP-CtIP to DSB-containing tracks in U2OS cells by laser micro-irradiation (21). An inhibition in the accumulation of GFP-CtIP to the laser tracks was observed upon depletion of PHF2, whereas the recruitment of mCherry-Nbs1 was unaffected in these conditions (Fig. 2A). To substantiate the effect of PHF2 on DNA end resection, the phosphorylation of RPA after chromatin fractionation was examined. These results confirmed the inhibition of IR-induced RPA phosphorylation after downregulation of PHF2 demonstrated earlier (Fig. 2B). This experiment additionally demonstrated that the IR-induced accumulation of HR protein Rad51 on the chromatin in control depleted cells was prevented upon knock down of PHF2 (Fig. 2B). In accordance, when analysing Rad51 focus formation in response to IR by immunofluorescence, we observed that knock down of PHF2, as well as depletion of CtIP, inhibited focus formation by Rad51 (Figure 2C). Together these results indicate that PHF2 promotes the DSB response by regulating DNA resection and strongly suggest that the subsequent DSB repair is affected by PHF2 depletion.

**Figure 2.**
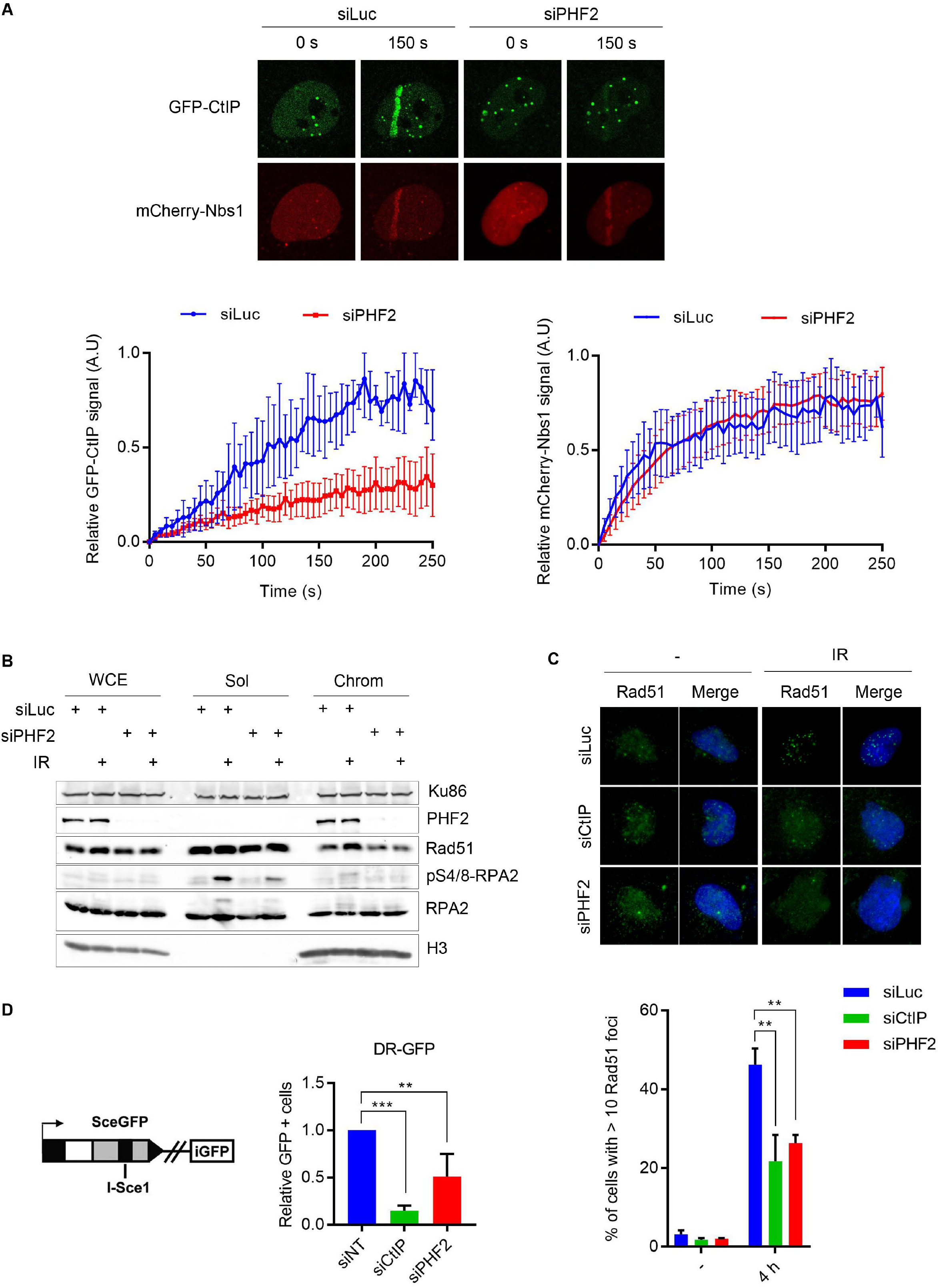
Depletion of PHF2 impairs DSB repair by homologous recombination. (**A**) U2OS cells, depleted for Luc or PHF2 by siRNA and 24 h later transfected with GFP-CtIP and mCherry-Nbs1, were laser-irradiated and analysed by time-lapse imaging. Upper panel: representative images of the indicated time points. Lower panel: relative fluorescence (left: GFP-CtIP, right: mCherry-Nbs1) at laser stripe in time of at least 50 cells per experiment. (**B**) U2OS cells were depleted for Luc or PHF2 by siRNA, treated with IR (10 Gy) and subjected to chromatin fractionation (WCE: whole cell extracts, Sol: soluble and Chrom: chromatin). Samples were analysed by western blot using the indicated antibodies. (**C**) U2OS cells were depleted for Luc, CtIP or PHF2 by siRNA. Cells were treated with IR (3 Gy), fixed after 4 h and Rad51 focus formation was analysed by immunofluorescence. Top panel: representative images. Bottom panel: quantification from three independent experiments with each at least 100 cells. (**D**) U2OS cells stably expressing a single copy of the DR-GFP reporter construct (see schematic overview) were depleted of CtIP, PHF2 or control (non-target, NT). After 48 hours, GFP fluorescence was analysed by flow cytometry. Presented is the relative fluorescence as compared to the control cells, of three independent experiments.

DNA end resection is a critical step in homology-directed DSB repair (4). To address whether PHF2 impacts on DSB repair by homologous recombination, we used a DR-GFP reporter assay in U2OS. In these cells, depletion of PHF2 caused a significant decrease in HR efficiency (Fig. 2D). The efficiency of Single Strand Annealing (SSA), another form of homology-directed repair that also depends on CtIP-mediated DNA end resection but is Rad51-independent (34), measured using an SA-GFP reporter assay, was also dramatically affected by PHF2 knock down (Fig. S1D). As HR depends on the availability of a sister chromatid as repair template, this type of repair is restricted to the S and G2 phases of the cell cycle (35). Importantly, PHF2 depletion does not affect the cell cycle distribution, as shown in figure S1E, demonstrating that the decrease in homology-directed DSB repair efficiency upon PHF2 knock down is not an indirect effect of changing the cell cycle.

### PHF2 directly controls CtIP and BRCA1 mRNA levels

We next set out to address if PHF2 affects DSB repair in a direct manner by acting at sites of DNA damage. To this end, the accumulation of Flag-PHF2 at sites of DNA damages generated by laser micro-irradiation was examined. Although mCherry-Nbs1 and γH2AX were detected at damaged regions upon laser micro-irradiation, we did not observe detectable Flag-PHF2 accumulation to such laser-tracks (Fig. S2A). In addition, also no accumulation of Flag-PHF2 was observed to a DSB created in a single genomic locus containing an array of LacO repeats following expression and tethering of a mCherry-LacI-FokI nuclease fusion (Fig. S2B). Together these results suggest that PHF2 regulates homology-directed DSB repair in an indirect and more global manner.

Interestingly, PHF2 contains a zinc finger-like PHD (plant homeodomain) finger, a motif found in proteins that are involved in transcriptional regulation, possibly by recognizing chromatin modifications (36), which suggests that PHF2 might regulate HR in a transcriptional manner. Western blot analysis of the levels of proteins involved in DSB repair demonstrated that downregulation of PHF2 led to diminished protein levels of CtIP as well as BRCA1, while leaving Rad51, RPA2, 53BP1, Nbs1 and Mre11 unaffected (Fig. 3A). Downregulating PHF2 by two additional siRNA oligonucleotides resulted in the same phenotype as seen before, namely a diminished abundance of CtIP and BRCA1 protein (Fig. 3B and C). In contrast, overexpression of Flag-PHF2 had the opposite effect: both BRCA1 and CtIP protein levels increased under these conditions (Fig. 3D). Furthermore, the lower protein levels of BRCA1 and CtIP upon depletion of PHF2 were partially rescued by expressing Flag-PHF2 (Fig. 3E). In addition, we could complement the effects of PHF2 depletion on focus formation of RPA2, Rad51 and 53BP1 by expressing an siRNA-resistant version of Flag-PHF2 (Fig. 3F, 3G and S1C, respectively). Together these data demonstrate that the effects of modulating PHF2 levels are genuinely due to depletion the of PHF2 instead of an off-target effect of the siRNA oligonucleotides used.

**Figure 3.**
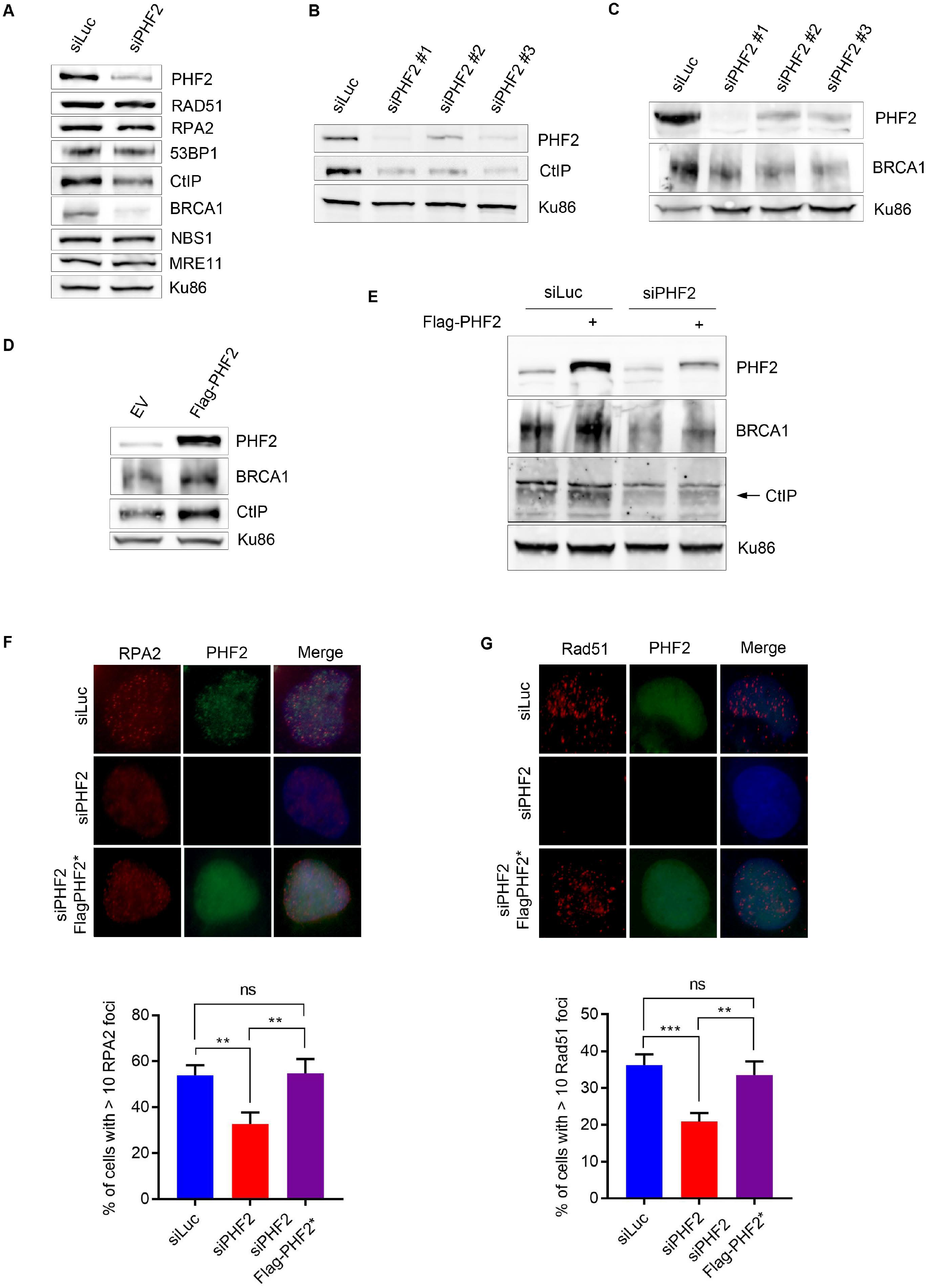
PHF2 regulates HR by modulating CtIP and BRCA1 levels. (**A**) U2OS cells were depleted for Luc or PHF2 by siRNA. 48 h later the cells were lysed and extracts were analysed by western blot with the indicated antibodies. (**B**) U2OS cells depleted for PHF2 using three different siRNA oligonucleotides. Western blot analysis using the indicated antibodies. (**C**) as in (B). (**D**) U2OS cells were transfected with empty vector (EV) or a Flag-PHF2 expression vector, followed by western blot analysis with the indicated antibodies. (**E**) U2OS cells were depleted for Luc or PHF2 by siRNA and 24 h later transfected with EV or Flag-PHF2. The day after, extracts were prepared and analysed by western blot with the indicated antibodies. (**F**) U2OS cells were depleted for PHF2 and transfected with siRNA-resistant Flag-PHF2 (Flag-PHF2*) the day after. One day later, cells were treated with IR (3 Gy) and fixed for IF after 1 hour. RPA2 focus formation of Flag-positive cells was analysed by immunofluorescence. Top panel: representative images. Bottom panel: quantification of three independent experiments with each at least 100 cells. (**G**) U2OS cells were depleted for PHF2 and transfected with siRNA-resistant Flag-PHF2 (Flag-PHF2*) the day after. One day later, cells were treated with IR (3 Gy) and fixed for IF after 4 hours. Rad51 focus formation of Flag-positive cells was analysed as in (F).

We next examined if PHF2 regulates CtIP and BRCA1 at the transcriptional level by investigating the effect of PHF2 depletion on *CtIP* and *BRCA1* mRNA in U2OS by quantitative PCR. The mRNA levels of *CtIP* and *BRCA1*, but not those of *Mre11* and *RPA2*, were decreased under conditions of PHF2 depletion when compared to control transfected cells (Fig. 4A), indicating that PHF2 controls homology-directed DSB repair by regulating the transcription of CtIP and BRCA1, two proteins critical for DNA end-resection and thereby initiation of HR.

**Figure 4.**
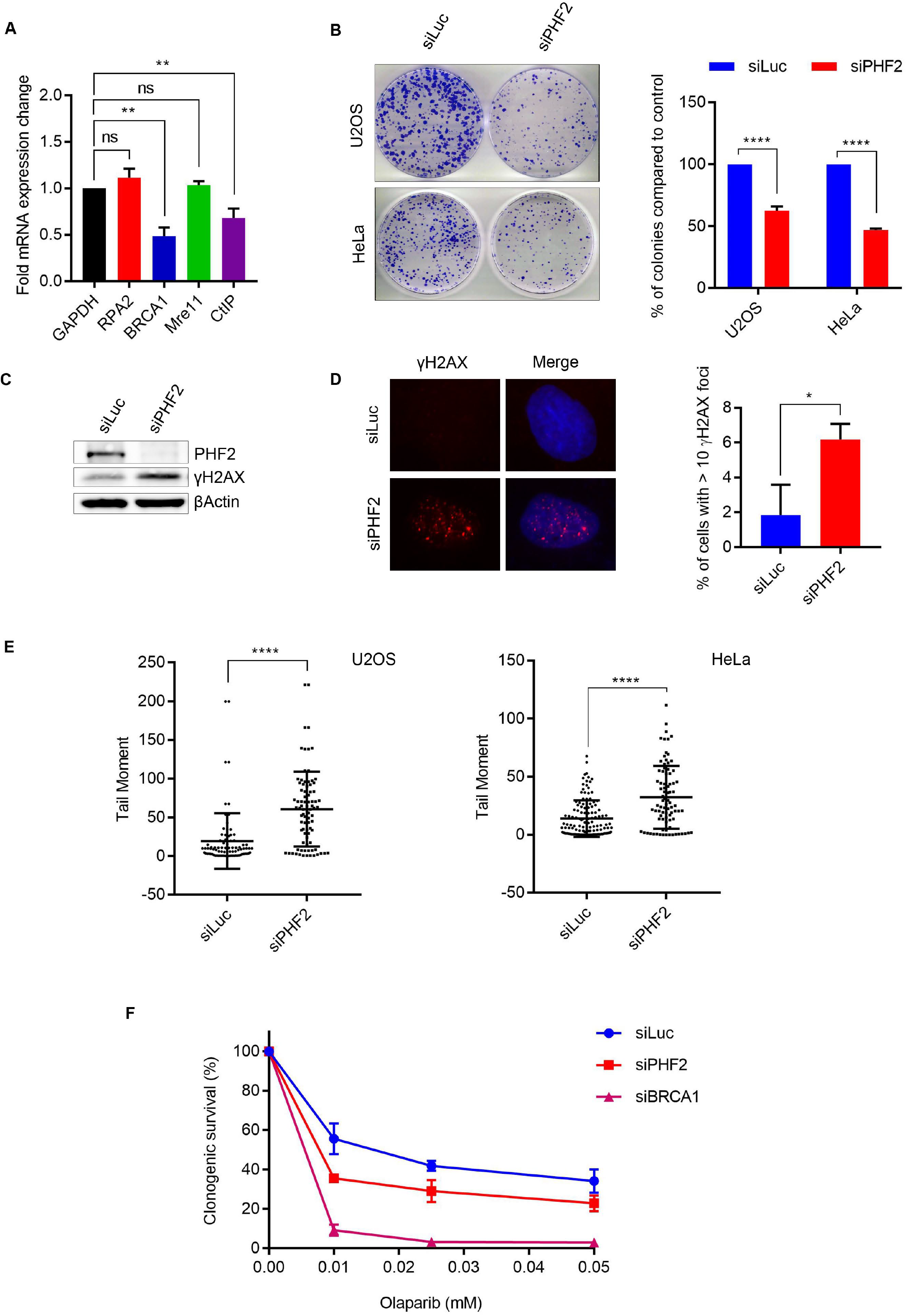
PHF2 controls the expression of CtIP and BRCA1 and prevents genome instability. (**A**) U2OS were downregulated for Luc or PHF2 by siRNA. 72 h later, RNA was isolated and mRNA levels of GAPDH, RPA2, BRCA1, Mre11 and CtIP were determined by RT-qPCR. Shown is the fold mRNA change of PHF2-depleted samples as compared to the Luc control. (**B**) U2OS and HeLa cells were depleted for PHF2 by siRNA. Equal numbers of cells were seeded and incubated for 10 days for colonies to form, after which the cells were fixed and stained (left panel). Quantification of the number of colonies from three independent experiments (right panel). (**C**) U2OS cells were depleted for Luc or PHF2 by siRNA. 48 h later the cells were lysed and analysed by western blot with the indicated antibodies. (**D**) As in (C), but now γH2AX focus formation was analysed by IF. Left panel: representative images. Right panel: quantification of three independent experiments with each at least 100 cells. (**E**) U2OS and HeLa cells, depleted for PHF2 by siRNA, were subjected to SCGE (see materials and methods). Depicted is the tail moment of three independent experiments. (**F**) HeLa cells were depleted for Luc, PHF2 or BRCA1 by siRNA. Cells were seeded and incubated for colony formation, in presence or absence of different concentrations of Olaparib.

### PHF2 downregulation causes genome instability and sensitivity to PARP inhibition

As HR is not only important for DSB repair but also plays a role in several other processes in unperturbed cells, the defective HR caused by downregulation of PHF2 is likely to affect genomic stability and cell survival in the absence of exogenous damage. Consistent with such a scenario, the colony forming capacity of both U2OS and HeLa cells decreased dramatically after depletion of PHF2 as compared to Luciferase knock down control cells (Fig. 4B). Western blot analysis demonstrated an increase in H2AX phosphorylation (γH2AX) in PHF2-depleted U2OS cells in the absence of exogenous damage, and this was confirmed by an elevated percentage of cells with γH2AX foci upon PHF2 knock down as compared to Luc control cells by immunofluorescence (Fig. 4C and D, respectively). To directly asses the appearance of DNA breaks resulting from downregulation of PHF2 in the absence of exogenous DNA damage in individual cells, we employed the alkaline comet assay. Compared to control depleted cells, decreasing PHF2 by siRNA led to a significant tail moment increase in both U2OS and Hela cells (Fig. 4E). Together these data demonstrate that regulated levels of PHF2 are important for the maintenance of genome stability.

Notably, HR deficiency is exploited in the treatment of tumours with BRCA1/BRCA2 mutations as cells in such conditions are sensitive to inhibition of poly(ADP-ribose) polymerase (PARP) (37). As PHF2 depletion affects HR, downregulation of this protein is likewise expected to result in sensitivity to PARP inhibition. Indeed, as knock down of BRCA1, depletion of PHF2 sensitized HeLa cells, although to a lesser extent, to inhibition of PARP1/2 by Olaparib, underscoring the importance of PHF2 in HR (Fig. 4F).

## DISCUSSION

In this study we show that the demethylase PHF2 contributes to the maintenance of genome integrity by controlling homology-directed DNA repair. The depletion of PHF2 thereby affects cell growth in unperturbed cells and the response to DNA damage. Specifically, we showed that downregulation of PHF2 affects the DNA damage-induced focus formation by BRCA1 and 53BP1, which together determine the choice between DSB repair by homologous recombination or non-homologous end joining. Concomitant with a decrease in BRCA1 focus formation, PHF2 depletion also affected the accumulation of CtIP to sites of DNA lesions. CtIP is known to initiate DNA end resection to generate ssDNA ends that are a prerequisite for HR (10, 21). BRCA1, that forms a complex with CtIP, collaborates in ssDNA-end formation (34, 38, 39). Indeed, impaired DSB resection was observed after PHF2 downregulation, as demonstrated by decreased RPA phosphorylation and focus formation. Consequently, PHF2 depletion impaired IR-induced focus formation of Rad51 and resulted in a diminished efficiency of HR. Impaired DSB repair by HR in response to PHF2 depletion is explained by our data showing that PHF2 controls the DSB response in a transcriptional manner, thereby leading to a decrease in *CtIP* and *BRCA1* mRNA and protein levels. That PHF2 regulates expression of CtIP and BRCA1 simultaneously is in accordance to indications in the literature that BRCA1 supports the control of CtIP. CtIP was reported to control its own transcription, possibly via interaction with BRCA1 through its BRCT domains (40, 41). BRCA1-dependent ubiquitination of CtIP is additionally required for CtIP chromatin binding and damage-induced focus formation (42), which likely explains the reported defect in the accumulation of GFP-CtIP at interstrand crosslinks induced by micro-irradiation (43). Our observation that depletion of PHF2 inhibits the accumulation of GFP-CtIP to the laser damage (Fig. 2A) is therefore likely to be an indirect effect of the diminished BRCA1 levels in these conditions. However, we cannot exclude the possibility that PHF2 regulates CtIP on several levels, like the recently reported splicing complex SF3B, that controls CtIP both at the level of mRNA abundance and the recruitment of CtIP to the chromatin in response to DNA damage (44).

The KDM7 family of histone demethylases, to which PHF2 belongs, can remove the methylation from H3K9, H3K27, or H4K20, which are responsible for transcriptional repression, by interaction of their PHD domain with methylated lysine residues and the subsequent removal of these methyl groups via the JmjC-domain (36, 45). Indeed, PHF2 was described to regulate transcription by removing the dimethylated H3K9 and, to a lesser extent, trimethylated H4K20 (46–49). We hypothesize that PHF2 controls *CtIP* and *BRCA1* gene transcription by erasing the transcription repression mark from the respective promoters (49). PHF2 was shown to stimulate the expression of genes driven by the transcription factors HNF4, CEBPα, p53 and NF-κB (46–49). Interestingly, NF-κB was shown to regulate HR by controlling BRCA1-CtIP complexes, although this effect is not mediated by transcriptional regulation of BRCA1 and CtIP (50). While we consider it likely that PHF2 demethylase activity is required for this function, we cannot rule out another mechanism for PHF2 action, as PHF2 was also demonstrated to repress transcription in a manner dependent on its methylated H3K4 binding activity yet independent on its demethylase activity (51). Instead, PHF2 was thought to repress transcription by competition with PHF8 for binding of the promoter and by recruiting SUV39H1, the H3K9me2/3 methyltransferase (51). In this respect, it is also important to mention that dimethylation of H4K20 is critical in the recruitment of 53BP1 to sites of DNA lesions (52, 53). Interfering with this methyl mark and/or competition for binding to methylated H4K20 by PHF2 could also contribute to the phenotype in disturbing homology-directed DSB repair observed in this study, although we consider this possibility less likely since we have no indications that PHF2 might function directly at the sites of DNA lesions.

By controlling homology-directed DSB repair, PHF2 emerges as a putative important regulator in maintaining genomic integrity, which is underscored by our results showing that knock down of PHF2 triggers DNA lesions in the absence of exogenous damage. Therefore, our data might have an importance in pathologies in which PHF2 levels are changed or PHF2 is mutated. For example, high PHF2 levels were found in oesophageal carcinoma and renal cell carcinoma (54, 55), whereas the PHF2 gene is deleted or hypermethylated in its promoter region in breast cancer (56). In addition, PHF2 mutants were reported in gastric and colon cancer (57). Together these observations suggest that PHF2 plays a role in the development and/or progression of cancer. Interestingly, the use of histone demethylases as therapeutic targets by pharmacological inhibitors is currently being investigated and might open new strategies for tumour therapy (58, 59). Our results demonstrating that cells depleted for PHF2 are more sensitive to inhibition of PARP might suggest that targeting PHF2 could be particularly effective in breast and ovarian cancers without mutations in BRCA1/2 or other known HR proteins.

## ACKNOWLEDGEMENTS

We thank Drs. Daniel Durocher and Xiaobing Shi for sharing reagents.

## REFERENCES

1. Bartek,J., Bartkova,J. and Lukas,J. (2007) DNA damage signalling guards against activated oncogenes and tumour progression. Oncogene, 26, 7773–7779.

2. Aguilera,A. and Gómez-González,B. (2008) Genome instability: a mechanistic view of its causes and consequences. Nat. Rev. Genet., 9, 204–217.

3. Hiom,K. (2010) Coping with DNA double strand breaks. DNA Repair (Amst.), 9, 1256–1263.

4. Huertas,P. (2010) DNA resection in eukaryotes: deciding how to fix the break. Nat. Struct. Mol. Biol., 17, 11–16.

5. Noordermeer,S.M., Adam,S., Setiaputra,D., Barazas,M., Pettitt,S.J., Ling,A.K., Olivieri,M., Álvarez-Quilón,A., Moatti,N., Zimmermann,M., et al. (2018) The shieldin complex mediates 53BP1-dependent DNA repair. Nature, 560, 117–121.

6. Mirman,Z., Lottersberger,F., Takai,H., Kibe,T., Gong,Y., Takai,K., Bianchi,A., Zimmermann,M., Durocher,D. and de Lange,T. (2018) 53BP1-RIF1-shieldin counteracts DSB resection through CST- and Polα-dependent fill-in. Nature, 560, 112–116.

7. Bunting,S.F., Callén,E., Wong,N., Chen,H.-T., Polato,F., Gunn,A., Bothmer,A., Feldhahn,N., Fernandez-Capetillo,O., Cao,L., et al. (2010) 53BP1 inhibits homologous recombination in Brca1-deficient cells by blocking resection of DNA breaks. Cell, 141, 243–254.

8. Chapman,J.R., Taylor,M.R.G. and Boulton,S.J. (2012) Playing the end game: DNA double-strand break repair pathway choice. Mol. Cell, 47, 497–510.

9. Symington,L.S. (2016) Mechanism and regulation of DNA end resection in eukaryotes. Crit. Rev. Biochem. Mol. Biol., 51, 195–212.

10. Jasin,M. and Rothstein,R. (2013) Repair of strand breaks by homologous recombination. Cold Spring Harb Perspect Biol, 5, a012740–a012740.

11. Sulli,G., Di Micco,R. and d’Adda di Fagagna,F. (2012) Crosstalk between chromatin state and DNA damage response in cellular senescence and cancer. Nature Publishing Group, 12, 709–720.

12. Burma,S., Chen,B.P., Murphy,M., Kurimasa,A. and Chen,D.J. (2001) ATM phosphorylates histone H2AX in response to DNA double-strand breaks. J. Biol. Chem., 276, 42462–42467.

13. Kolas,N.K., Chapman,J.R., Nakada,S., Ylanko,J., Chahwan,R., Sweeney,F.D., Panier,S., Mendez,M., Wildenhain,J., Thomson,T.M., et al. (2007) Orchestration of the DNA-Damage Response by the RNF8 Ubiquitin Ligase. Science, 318, 1637–1640.

14. Doil,C., Mailand,N., Bekker-Jensen,S., Menard,P., Larsen,D.H., Pepperkok,R., Ellenberg,J., Panier,S., Durocher,D., Bartek,J., et al. (2009) RNF168 Binds and Amplifies Ubiquitin Conjugates on Damaged Chromosomes to Allow Accumulation of Repair Proteins. Cell, 136, 435–446.

15. Stewart,G.S., Panier,S., Townsend,K., Al-Hakim,A.K., Kolas,N.K., Miller,E.S., Nakada,S., Ylanko,J., Olivarius,S., Mendez,M., et al. (2009) The RIDDLE Syndrome Protein Mediates a Ubiquitin-Dependent Signaling Cascade at Sites of DNA Damage. Cell, 136, 420–434.

16. Zhao,Y., Brickner,J.R., Majid,M.C. and Mosammaparast,N. (2014) Crosstalk between ubiquitin and other post-translational modifications on chromatin during double-strand break repair. Trends Cell Biol., 24, 426–434.

17. Kouzarides,T. (2000) Acetylation: a regulatory modification to rival phosphorylation? EMBO J., 19, 1176–1179.

18. Greer,E.L. and Shi,Y. (2012) Histone methylation: a dynamic mark in health, disease and inheritance. Nat. Rev. Genet., 13, 343–357.

19. Luijsterburg,M.S. and van Attikum,H. (2011) Chromatin and the DNA damage response: the cancer connection. Mol Oncol, 5, 349–367.

20. Rabl,J., Bunker,R.D., Schenk,A.D., Cavadini,S., Gill,M.E., Abdulrahman,W., Andrés-Pons,A., Luijsterburg,M.S., Ibrahim,A.F.M., Branigan,E., et al. (2019) Structural Basis of BRCC36 Function in DNA Repair and Immune Regulation. Mol. Cell, 75, 483–497.e9.

21. Sartori,A.A., Lukas,C., Coates,J., Mistrik,M., Fu,S., Bartek,J., Baer,R., Lukas,J. and Jackson,S.P. (2007) Human CtIP promotes DNA end resection. Nature, 450, 509–514.

22. Luijsterburg,M.S., de Krijger,I., Wiegant,W.W., Shah,R.G., Smeenk,G., de Groot,A.J.L., Pines,A., Vertegaal,A.C.O., Jacobs,J.J.L., Shah,G.M., et al. (2016) PARP1 Links CHD2-Mediated Chromatin Expansion and H3.3 Deposition to DNA Repair by Non-homologous End-Joining. Mol. Cell, 61, 547–562.

23. Pierce,A.J., Johnson,R.D., Thompson,L.H. and Jasin,M. (1999) XRCC3 promotes homology-directed repair of DNA damage in mammalian cells. Genes Dev., 13, 2633–2638.

24. Bennardo,N., Cheng,A., Huang,N. and Stark,J.M. (2008) Alternative-NHEJ is a mechanistically distinct pathway of mammalian chromosome break repair. PLoS Genet., 4, e1000110.

25. Méndez,J. and Stillman,B. (2000) Chromatin association of human origin recognition complex, cdc6, and minichromosome maintenance proteins during the cell cycle: assembly of prereplication complexes in late mitosis. Mol. Cell. Biol., 20, 8602–8612.

26. Smits,V.A.J., Reaper,P.M. and Jackson,S.P. (2006) Rapid PIKK-dependent release of Chk1 from chromatin promotes the DNA-damage checkpoint response. Curr. Biol., 16, 150–159.

27. Typas,D., Luijsterburg,M.S., Wiegant,W.W., Diakatou,M., Helfricht,A., Thijssen,P.E., van den Broek,B., van de Broek,B., Mullenders,L.H. and van Attikum,H. (2015) The de-ubiquitylating enzymes USP26 and USP37 regulate homologous recombination by counteracting RAP80. Nucleic Acids Res., 43, 6919–6933.

28. Tang,J., Cho,N.W., Cui,G., Manion,E.M., Shanbhag,N.M., Botuyan,M.V., Mer,G. and Greenberg,R.A. (2013) Acetylation limits 53BP1 association with damaged chromatin to promote homologous recombination. Nat. Struct. Mol. Biol., 20, 317–325.

29. Hu,Y., Scully,R., Sobhian,B., Xie,A., Shestakova,E. and Livingston,D.M. (2011) RAP80-directed tuning of BRCA1 homologous recombination function at ionizing radiation-induced nuclear foci. Genes Dev., 25, 685–700.

30. Smits,V.A.J. and Gillespie,D.A. (2015) DNA damage control: regulation and functions of checkpoint kinase 1. FEBS J., 282, 3681–3692.

31. Zou,L. and Elledge,S.J. (2003) Sensing DNA damage through ATRIP recognition of RPA-ssDNA complexes. Science, 300, 1542–1548.

32. Shiotani,B. and Zou,L. (2009) Single-stranded DNA orchestrates an ATM-to-ATR switch at DNA breaks. Mol. Cell, 33, 547–558.

33. Maréchal,A. and Zou,L. (2015) RPA-coated single-stranded DNA as a platform for post-translational modifications in the DNA damage response. Cell Res., 25, 9–23.

34. Yun,M.H. and Hiom,K. (2009) CtIP-BRCA1 modulates the choice of DNA double-strand-break repair pathway throughout the cell cycle. Nature, 459, 460–463.

35. Huertas,P. and Jackson,S.P. (2009) Human CtIP mediates cell cycle control of DNA end resection and double strand break repair. J. Biol. Chem., 284, 9558–9565.

36. Fortschegger,K. and Shiekhattar,R. (2011) Plant homeodomain fingers form a helping hand for transcription. Epigenetics, 6, 4–8.

37. Jackson,S.P. and Helleday,T. (2016) DNA REPAIR. Drugging DNA repair. Science, 352, 1178–1179.

38. Chen,L., Nievera,C.J., Lee,A.Y.-L. and Wu,X. (2008) Cell cycle-dependent complex formation of BRCA1.CtIP.MRN is important for DNA double-strand break repair. J. Biol. Chem., 283, 7713–7720.

39. Nakamura,K., Kogame,T., Oshiumi,H., Shinohara,A., Sumitomo,Y., Agama,K., Pommier,Y., Tsutsui,K.M., Tsutsui,K., Hartsuiker,E., et al. (2010) Collaborative action of Brca1 and CtIP in elimination of covalent modifications from double-strand breaks to facilitate subsequent break repair. PLoS Genet., 6, e1000828.

40. Yu,X., Wu,L.C., Bowcock,A.M., Aronheim,A. and Baer,R. (1998) The C-terminal (BRCT) domains of BRCA1 interact in vivo with CtIP, a protein implicated in the CtBP pathway of transcriptional repression. J. Biol. Chem., 273, 25388–25392.

41. Liu,F. and Lee,W.-H. (2006) CtIP activates its own and cyclin D1 promoters via the E2F/RB pathway during G1/S progression. Mol. Cell. Biol., 26, 3124–3134.

42. Yu,X., Fu,S., Lai,M., Baer,R. and Chen,J. (2006) BRCA1 ubiquitinates its phosphorylation-dependent binding partner CtIP. Genes Dev., 20, 1721–1726.

43. Duquette,M.L., Zhu,Q., Taylor,E.R., Tsay,A.J., Shi,L.Z., Berns,M.W. and McGowan,C.H. (2012) CtIP is required to initiate replication-dependent interstrand crosslink repair. PLoS Genet., 8, e1003050.

44. Prados-Carvajal,R., López-Saavedra,A., Cepeda-García,C., Jimeno,S. and Huertas,P. (2018) Multiple roles of the splicing complex SF3B in DNA end resection and homologous recombination. DNA Repair (Amst.), 66-67, 11–23.

45. Jaskelioff,M. and Peterson,C.L. (2003) Chromatin and transcription: histones continue to make their marks. In. Nature Publishing Group, Vol. 5, pp. 395–399.

46. Baba,A., Ohtake,F., Okuno,Y., Yokota,K., Okada,M., Imai,Y., Ni,M., Meyer,C.A., Igarashi,K., Kanno,J., et al. (2011) PKA-dependent regulation of the histone lysine demethylase complex PHF2-ARID5B. Nat. Cell Biol., 13, 668–675.

47. Stender,J.D., Pascual,G., Liu,W., Kaikkonen,M.U., Do,K., Spann,N.J., Boutros,M., Perrimon,N., Rosenfeld,M.G. and Glass,C.K. (2012) Control of proinflammatory gene programs by regulated trimethylation and demethylation of histone H4K20. Mol. Cell, 48, 28–38.

48. Okuno,Y., Ohtake,F., Igarashi,K., Kanno,J., Matsumoto,T., Takada,I., Kato,S. and Imai,Y. (2013) Epigenetic regulation of adipogenesis by PHF2 histone demethylase. Diabetes, 62, 1426–1434.

49. Lee,K.-H., Park,J.-W., Sung,H.-S., Choi,Y.-J., Kim,W.H., Lee,H.S., Chung,H.-J., Shin,H.-W., Cho,C.-H., Kim,T.-Y., et al. (2015) PHF2 histone demethylase acts as a tumor suppressor in association with p53 in cancer. Oncogene, 34, 2897–2909.

50. Volcic,M., Karl,S., Baumann,B., Salles,D., Daniel,P., Fulda,S. and Wiesmüller,L. (2012) NF-κB regulates DNA double-strand break repair in conjunction with BRCA1-CtIP complexes. Nucleic Acids Res., 40, 181–195.

51. Shi,G., Wu,M., Fang,L., Yu,F., Cheng,S., Li,J., Du,J.X. and Wong,J. (2014) PHD finger protein 2 (PHF2) represses ribosomal RNA gene transcription by antagonizing PHF finger protein 8 (PHF8) and recruiting methyltransferase SUV39H1. Journal of Biological Chemistry, 289, 29691–29700.

52. Botuyan,M.V., Lee,J., Ward,I.M., Kim,J.-E., Thompson,J.R., Chen,J. and Mer,G. (2006) Structural Basis for the Methylation State-Specific Recognition of Histone H4-K20 by 53BP1 and Crb2 in DNA Repair. Cell, 127, 1361–1373.

53. Pei,H., Zhang,L., Luo,K., Qin,Y., Chesi,M., Fei,F., Bergsagel,P.L., Wang,L., You,Z. and Lou,Z. (2011) MMSET regulates histone H4K20 methylation and 53BP1 accumulation at DNA damage sites. Nature, 469, 124–128.

54. Sun,L.-L., Sun,X.-X., Xu,X.-E., Zhu,M.-X., Wu,Z.-Y., Shen,J.-H., Wu,J.-Y., Huang,Q., Li,E.-M. and Xu,L.-Y. (2013) Overexpression of Jumonji AT-rich interactive domain 1B and PHD finger protein 2 is involved in the progression of esophageal squamous cell carcinoma. Acta Histochem., 115, 56–62.

55. Lee,C., Kim,B., Song,B. and Moon,K.C. (2017) Implication of PHF2 Expression in Clear Cell Renal Cell Carcinoma. J Pathol Transl Med, 51, 359–364.

56. Sinha,S., Singh,R.K., Alam,N., Roy,A., Roychoudhury,S. and Panda,C.K. (2008) Alterations in candidate genes PHF2, FANCC, PTCH1 and XPA at chromosomal 9q22.3 region: pathological significance in early- and late-onset breast carcinoma. Mol. Cancer, 7, 84.

57. Lee,J.H., Yoo,N.J., Kim,M.S. and Lee,S.H. (2017) Histone Demethylase Gene PHF2 Is Mutated in Gastric and Colorectal Cancers. Pathol. Oncol. Res., 23, 471–476.

58. Højfeldt,J.W., Agger,K. and Helin,K. (2013) Histone lysine demethylases as targets for anticancer therapy. Nat Rev Drug Discov, 12, 917–930.

59. Morera,L., Lübbert,M. and Jung,M. (2016) Targeting histone methyltransferases and demethylases in clinical trials for cancer therapy. Clin Epigenetics, 8, 57.

