## Supplemental data for "PHF2 regulates homology-directed DNA repair by controlling the resection of DNA double strand breaks"

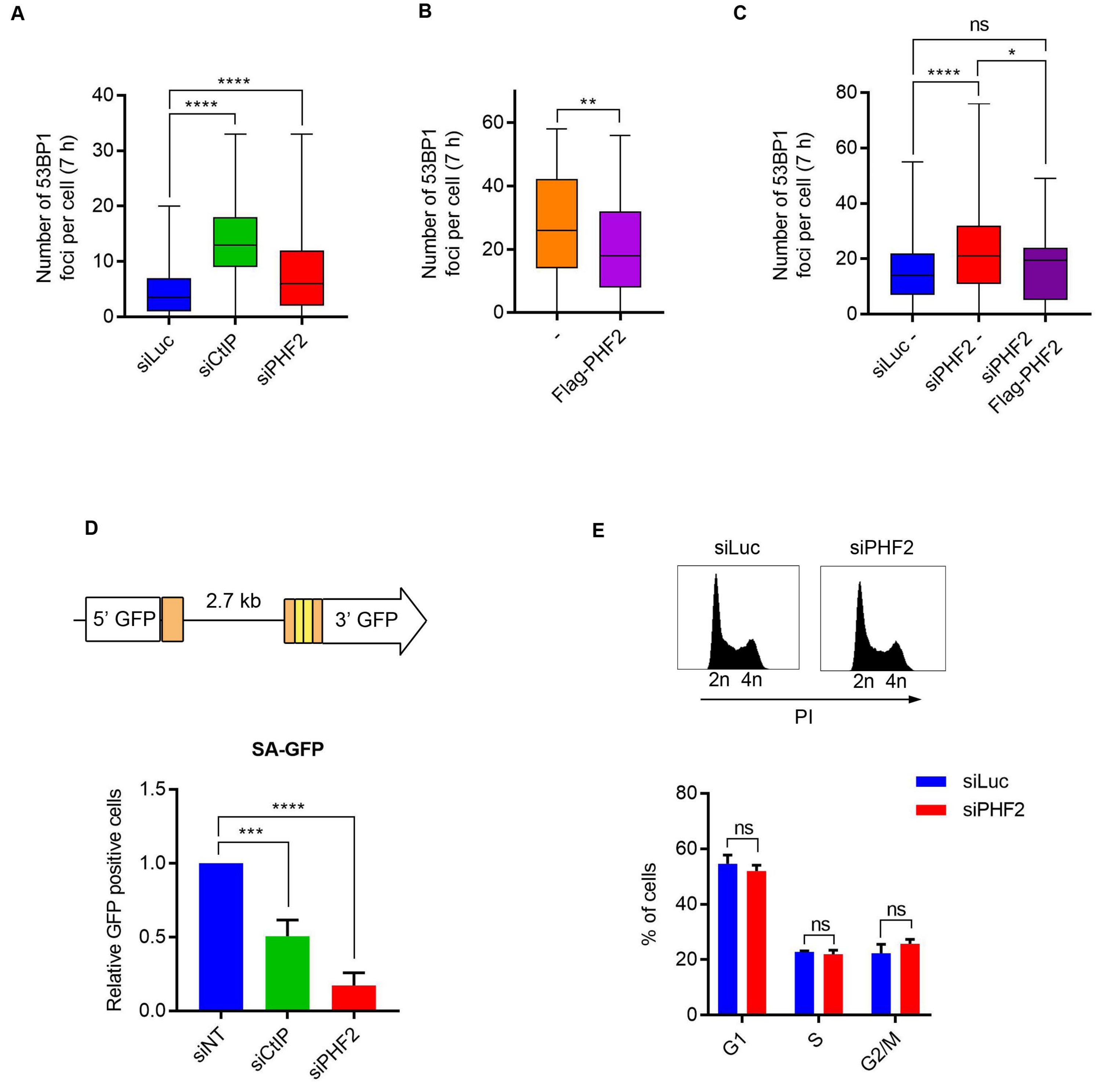

A
Number of 53BP1 foci per cell (7 h)
40
30
20
10
0
****
****
siLuc
siCtIP
siPHF2
B
Number of 53BP1 foci per cell (7 h)
60
40
20
0
**
-
Flag-PHF2
C
ns
*
****
Number of 53BP1 foci per cell (7 h)
80
60
40
20
0
siLuc -
siPHF2 -
siPHF2 Flag-PHF2
D
5' GFP
2.7 kb
3' GFP
SA-GFP
Relative GFP positive cells
1.5
1.0
0.5
0.0
***
****
siNT
siCtIP
siPHF2
E
siLuc
siPHF2
2n 4n
2n 4n
PI
siLuc
siPHF2
% of cells
80
60
40
20
0
ns
ns
ns
G1
S
G2/M


Figure S1

**A**

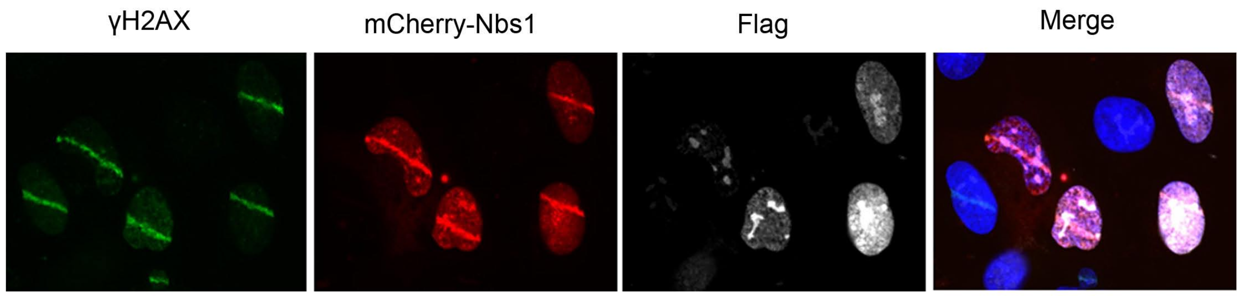


**B**

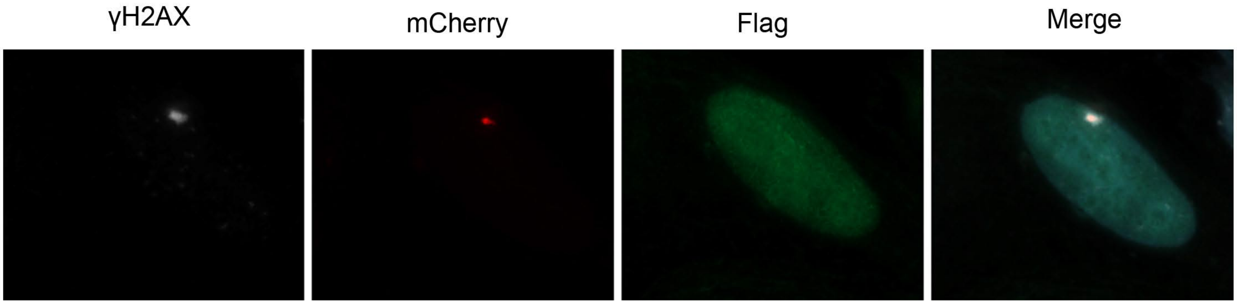


Figure S2

**Supplemental Figure Legends**

**Figure S1.** (**A**) U2OS cells were depleted for Luciferase (Luc), CtIP or PHF2 by siRNA. After 48 h, cells were treated with IR (10 Gy) and fixed after 7 hours. 53BP1 focus formation was analysed by immunofluorescence. Shown is the quantification of the number of 53BP1 foci per cell of three independent experiments with each at least 100 cells. (**B**) U2OS cells were transfected with Flag-PHF2, treated with 3 Gy IR, fixed 7 hours later and subsequently analysed by immunofluorescence for Flag and 53BP1. 53BP1 focus formation was analysed as in (A). (**C**) U2OS cells were depleted for PHF2 and transfected with siRNA-resistant Flag-PHF2 the day after. One day later, cells were treated with IR (3 Gy) and fixed for IF after 7 hours. 53BP1 focus formation was analysed as in (A). (**D**) U2OS cells carrying the SA-GFP reporter construct were depleted of CtIP, PHF2 or control (NT). After 48 hours, GFP fluorescence was analysed by FACS. Presented is the relative fluorescence as compared to the control cells, of three independent experiments. (**E**) U2OS cells were downregulated for Luc or PHF2 by siRNA. 48 hours later cells were fixed, stained with PI and analysed by flow cytometry. Quantifications shows the percentage of cells in G1, S or G2/M phases of three independent experiments.

**Figure S2.** (**A**) U2OS cells transfected with mCherry-Nbs1 and Flag-PHF2 were laser-irradiated and fixed, followed by immunofluorescence analysis for γH2AX and Flag. (**B**) U2OS 2-6-3 cells expressing inducible FokI-mCherry-LacR were transfected with Flag-PHF2 and treated to induce FokI. After fixation, cells were analysed by immunofluorescence using the indicated antibodies.
